# An Organotypic Mammary Duct Model Capturing Distinct Events of DCIS Progression

**DOI:** 10.1101/2020.08.06.240242

**Authors:** Jonathan Kulwatno, Xiangyu Gong, Rebecca DeVaux, Jason I. Herschkowitz, Kristen Lynn Mills

## Abstract

Ductal carcinoma *in situ* (DCIS) is a pre-cancerous stage breast cancer, where abnormal cells are contained within the duct, but have not invaded into the surrounding tissue. However, only 30-40% of DCIS cases are likely to progress into an invasive ductal carcinoma (IDC), while the remainder are innocuous. Since little is known about what contributes to the transition from DCIS to IDC, clinicians and patients tend to opt for treatment, leading to concerns of overdiagnosis and overtreatment. *In vitro* models are currently being used to probe how DCIS transitions into IDC, but many models do not take into consideration the macroscopic tissue architecture and the biomechanical properties of the microenvironment. Here, we developed an organotypic mammary duct model by molding a channel within a collagen matrix and lining it with a basement membrane. By adjusting the concentration of collagen, we effectively modulated the stiffness and morphological properties of the matrix and examined how an assortment of breast cells responded to changing density and stiffness of the matrix. We first validated the model using two established, phenotypically divergent breast cancer cell lines by demonstrating the ability of the cells to either invade (MDA-MB-231) or cluster (MCF7). We then examined how cells of the isogenic MCF10 series—spanning the range from healthy to aggressive—behaved within our model and observed distinct characteristics of breast cancer progression such as hyperplasia and invasion, in response to collagen concentration. Our results show that the model can recapitulate different stages of breast cancer progression and that the MCF10 series is adaptable to physiologically relevant *in vitro* studies, demonstrating the potential of both the model and cell lines to elucidate key factors that may contribute to understanding the transition from DCIS to IDC.

**IMPACT STATEMENT:** The success of early preventative measures for breast cancer has left patients susceptible to overdiagnosis and overtreatment. Limited knowledge of factors driving an invasive transition has inspired the development of *in vitro* models that accurately capture this phenomenon. However, current models tend to neglect the macroscopic architecture and biomechanical properties of the mammary duct. Here, we introduce an organotypic model that recapitulates the cylindrical geometry of the tissue and the altered stroma seen in tumor microenvironments. Our model was able to capture distinct features associated with breast cancer progression, demonstrating its potential to uncover novel insights into disease progression.

## INTRODUCTION

In the late 1980s female breast cancer diagnoses—still the most common female cancer—increased, but the number of associated deaths since then has continuously declined ^1^. This has been widely attributed to improvements in diagnostic technologies and strategies that enable clinicians to detect breast cancer development at its early stages^2–4^. A quarter of breast cancers are diagnosed as a ductal carcinoma *in situ* (DCIS) ^5^, which is considered a pre-cancerous stage with abnormal cells confined within the mammary duct. When diagnosed with DCIS, clinicians and patients tend to opt for treatments such as surgery and radiation to prevent transition to the malignant invasive ductal carcinoma (IDC). However, only 30-40% of DCIS cases are likely to progress into IDC, while the remainder are innocuous ^6,7^. Survivorship is as high as 98% ^5^, but the preemptive approach has raised concerns over the anxiety and negative impact on patients’ lives as well as healthcare costs associated with overdiagnosis and overtreatment ^2,3,8^. Without a better understanding of the factors that drive DCIS to transition to IDC, apprehensions about forgoing preemptive treatment will naturally remain.

In the quest to identify and validate targets related to the transition of DCIS to IDC, a multitude of models have been employed. Mouse models and two-dimensional cultures have contributed greatly, but concerns surrounding the ability to tease out confounding factors and physiological relevance, respectively, pervade. A promising approach to address those concerns is three-dimensional, *in vitro* models ^9,10^. In a recent review on *in vitro* models, Brock and colleagues conclude that 3D models of DCIS are more relevant for testing drug efficacy and toxicity in breast cancer due to the tissue-mimicking architecture, which provides a measure of the *in vivo* biological counterbalancing influences ^11^. Simply, 3D *in vitro* models allow for cellcell and cell-matrix interplay with an added dimension that better simulates the native environment. However, many 3D models are simply tumor cells or aggregates embedded in uniform matrices that do not account for tissue-scale architecture or the morphological and mechanical properties of the tissue, both of which affect cell organization and function.

A mammary duct is a tubular organ that can be simplified as a cell-lined cylinder within a matrix. Bischel and colleagues have highlighted the importance of 3D tubular models of mammary duct structure, finding that such models alter levels of secreted factors and cytokines in addition to the oft-observed organizational impact on the cells ^12^. Models of this sort have made use of microfluidics techniques to line cells on the lumen and influence their orientation and organization in a physiologically relevant manner ^11,13^. These models have allowed researchers to isolate and study the effects of stromal cells on breast cancer cell invasion, apoptosis, and estrogen sensitivity; in addition to metabolic adaptations due to hypoxia ^14–20^. However, these organotypic models have not yet been used to explore how the biomechanical properties of the microenvironment—in tandem with tissue architecture—contribute to disease progression.

The tumor microenvironment has been well recognized to often be denser and stiffer than its healthy counterpart due to an increase in collagen deposition and crosslinking via a desmoplastic response. The increased tissue stiffness and collagen density has been linked to an increasing malignant phenotype of mammary epithelial cells ^21–23^. This knowledge has launched a plethora of investigations that have tried to unravel the relationships between the mechanical microenvironment and/or the collagen density and structure with breast cancer behaviors ^24–29^. These types of experiments, however, have not used models representative of the architecture of the mammary duct.

In this study, we aimed to develop an organotypic mammary duct model that is architecturally relevant and with which we could explore the effects of collagen concentration on the organization and phenotype of normal mammary epithelial and breast cancer cells. A simple templating device was used to form a cylindrical structure within collagen gels of different concentrations—to recapitulate the architecture of the mammary duct at different clinical stages—and then lined with a basement membrane. We validated our model using two phenotypically divergent breast cancer cell lines (MCF7 and MDA-MB-231) and observed expected behaviors. The model was then used to study how isogenic cells of increasing malignancy (MCF10 series) responded to collagen concentration.

## MATERIALS & METHODS

### Cell culture

MDA-MB-231 human breast cancer cells were donated by Dr. Aram Chung (RPI). The cells were maintained as monocultures in RPMI-1640 (Gibco) supplemented with 5% FBS (Gibco), 2 mM L-glutamine, and 1x antimycotic/antibiotic solution (Gibco). MCF7 human breast cancer cells were purchased from ATCC. Cells were maintained as monocultures in DMEM (Corning) supplemented with 10% FBS (Gibco) and 1x antimycotic/antibiotic solution (Gibco). The MCF10A human breast epithelial cell line and its related series (MCF10DCIS.COM and MCF10CA1) were provided by Dr. Jason Herschkowitz (University at Albany). Cells were maintained as monocultures in DMEM/F-12 (Corning) supplemented with 5% horse serum (Gibco), 20 ng/mL EGF (ThermoFisher), 500 μg/mL hydrocortisone (Sigma), 100 ng/mL cholera toxin (Enzo Life Sciences), 10 μg/mL insulin (MP Biomedical), and 1x antimycotic/antibiotic solution (Gibco).

All cells were cultured in an incubator at 37 °C and 5% CO_2_. Upon reaching 80-90% confluency, cells were collected using 0.05% trypsin (Gibco), centrifuged to remove collection media, and replaced with fresh media.

### Rat tail collagen extraction

Sprague Dawley adult female rat tails were generously provided by Dr. Ryan Gilbert (RPI). For extraction, protocols similar to Ritte and Rajan *et al*. were used ^30,31^. Briefly, tail tendons were isolated, washed, and dissolved in acetic acid on a magnetic stir plate for 3 days at 4 °C. After centrifugation, the supernatant was collected, frozen at −80 °C, and then lyophilized with assistance from Dr. Mariah Hahn (RPI). The resulting collagen mesh was collected, weighed, and resuspended at 10 mg/mL in fresh 20 mM acetic acid in a glass vial and kept at 4 °C. The collagen was sterilized with chloroform. Purity was verified via SDS-PAGE and circular dichroism by comparison of a commercially available rat tail collagen solution (Invitrogen) (Supplemental Figure 1).

### Rheology

Shear modulus was determined using an AR-G2 (TA Instruments) as described in Zuidema *et al*.^32^. Briefly, 250 μL of cold collagen precursor solution, at the desired concentration, was pipetted onto the rheometer plate and a 20 mm diameter plate test geometry was lowered to a 500 μm gap. Samples were allowed to equilibrate at 37 °C for 45 minutes and then characterized via a time sweep at 1 Hz and 10% strain at 37 °C for 15 minutes. A solvent trap containing deionized water was used to prevent evaporation.

### Organotypic mammary duct fabrication

To generate the mammary duct models, a three-dimensional support with inner dimensions of 15 mm × 10 mm × 2.25 mm that contained 1 mm-diameter inlets on opposite ends (Figure 1A) was printed with acrylonitrile butadiene styrene (ABS) using a 3D printer (UPrint SE Plus, Stratasys). Polydimethylsiloxane (PDMS) was poured into the two larger pockets and cured to provide a sealing surface. A trimmed microscope slide (Glove Scientific) with dimensions of 26 mm × 25 mm was placed below the template and secured by binder clips that bonded the slide and the PDMS, ensuring a seal between the glass and the template. A blunt 20-gauge needle (20G; BD, O.D. = 0.908 mm) was inserted through both inlets, spanning the length of the support structure. All components were soaked in 70% ethanol overnight to prevent contamination.

**Figure 1.**
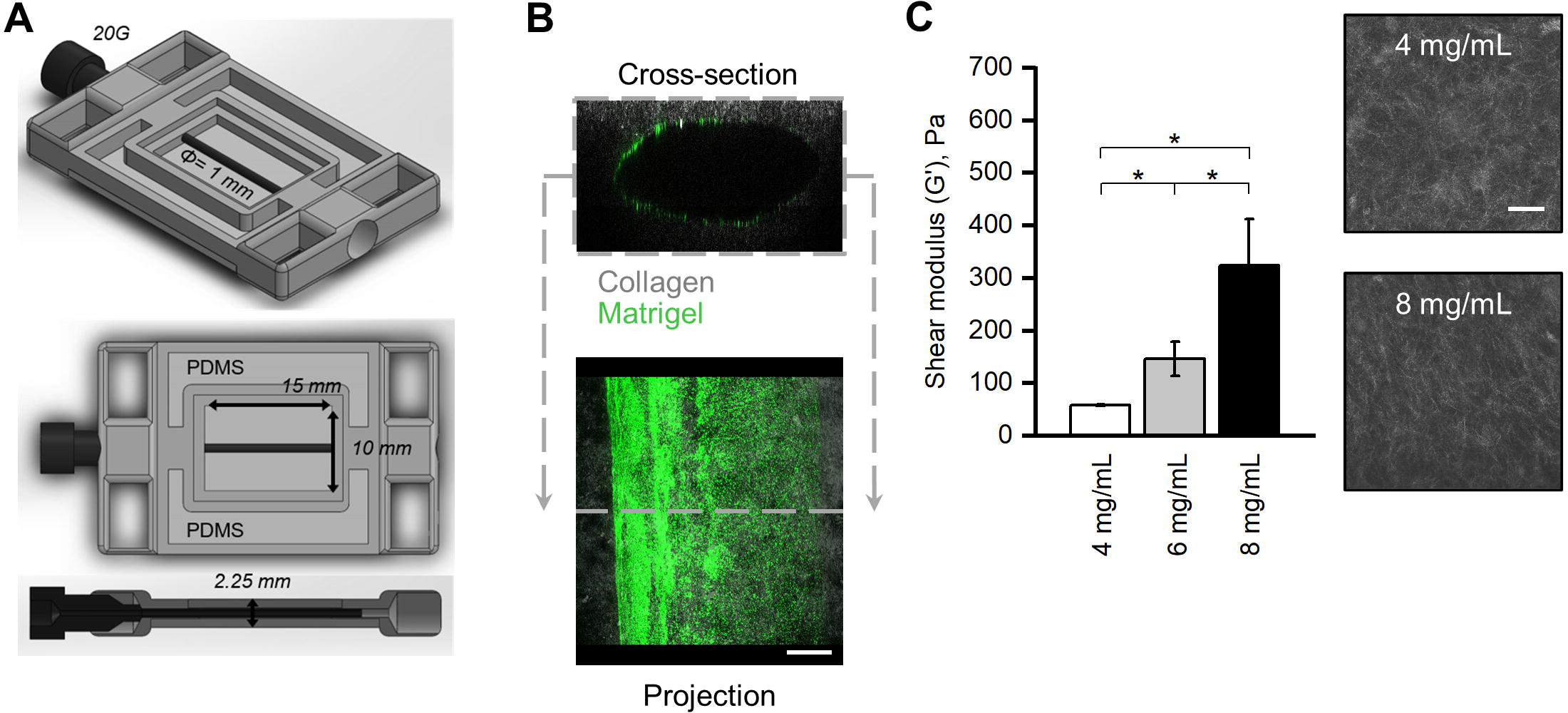
Organotypic mammary duct models with tunable mechanical and morphological properties. A) 3D printed support used to template a microchannel within a collagen matrix. B) The organotypic mammary duct model as a hollow channel within a collagen matrix (gray) lined with a basement membrane (green). Scale bar is 200 μm. C) Left: shear modulus of collagen gels of different concentrations. Data is presented as mean ± S.D. from at least three independent measurements. * denotes p < 0.05 via Student’s T-test. Right: representative images of collagen microstructure from collagen gels of different concentrations. Scale bar is 50 μm.

The 10 mg/mL collagen solution, described above, was neutralized by adding 1-part 10x PBS, 1-part NaOH and water to 8-parts collagen solution on ice. A final concentration of 8 mg/mL and a pH of 7.0 was confirmed. The solution was diluted to 4 mg/mL when necessary using cold media of the respective cell line. The desired collagen precursor solution was pipetted into the template (475 μL) and allowed to gel at 37 °C for 45 minutes, after which the needle was carefully removed. The constructs were cooled on ice for 5 minutes before the lumens were injected with a 2 mg/mL Matrigel solution (Corning) diluted with the respective media and containing fluorescence microbeads at 5 × 10^9^ beads/mL (ThermoFisher). The Matrigel solution was allowed to coat the collagen for 10 minutes before pipetting a cell suspension into the channel. For the validation studies, MDA-MB-231 and MCF7 cells were seeded at 250 cells/mm^2^ and 1200 cells/mm^2^, respectively. For the comparison study, MCF7, MCF10A, MCF10DCIS.COM, and MCF10CA1 cells were seeded at 5000 cells/mm^2^. Constructs were flipped after 15 minutes to ensure even coverage. Samples were then transferred into a 6-well plate and cultured in an incubator at 37 °C at 5% CO_2_ for 72-hours for the validation studies or 1 week for the comparison study.

### F-actin and nuclei staining

After the specified growth period, the mammary duct models were fixed with a 4% paraformaldehyde solution containing 0.01% glutaraldehyde for 30 minutes at room temperature while shaking. Samples were then washed with PBS three times for five minutes each before adding a solution containing 0.01% Triton X-100 in PBS with rhodamine phalloidin (1:1000; ThermoFisher) and Hoechst (1:10,000; ThermoFisher) overnight while shaking. Samples were again washed with and left in PBS.

### Image acquisition and analysis

A Leica SP8 was used to capture the fluorescence images of F-actin via rhodamine phalloidin and cell nuclei via Hoechst. Confocal reflectance microscopy was used to image collagen architecture. Images (10X objective) were taken bottom-up with the channel orthogonal to the objective and parallel to the stage. ImageJ was used to reconstruct, view, and analyze confocal slices. Observations of luminal surface coverage and organization of cells were made on maximum projection images of the z-stacks (Supplemental Figure 2B, image type 1). Measurements of cell morphology were done on individual slices of 2 μm steps within z-stacks. Invasion, nuclei distances, and F-actin coverage were done on leftward re-sliced cross-sections of the z-stacks at 20 μm steps (Supplemental Figure 2B, image type 2). Separate confocal slices were captured using a higher magnification objective (40X) at the mid-height plane of the channel at the boundary of the lumen and matrix to be able to directly measure the thickness of the cell layer and observe cell invasion into the matrix (Supplemental Figure 2B, image type 3).

### Statistics

Statistical analyses and data plotting were performed using OriginPro (OriginLab). Kruskal-Wallis ANOVA was used to compare within cell lines after normality was rejected using Kolmogorov-Smirnov tests. Box plots are presented with outliers for clarity of data visualization. Statistical differences were considered significant at p < 0.05.

## RESULTS

### Manufacturing of an organotypic mammary duct with tunable matrix mechanical and morphological properties

The mammary duct is a tubular organ within the connective tissue that can be simplified as a cylindrical channel within a matrix. Thus, we developed a 3D printed support that could hold a needle suspended over a surface, which would be used to template a channel within a matrix (Figure 1A). A 20 gauge needle with an outer diameter of 0.908 mm was used to closely match the average diameter of a human mammary duct ^33,34^. From there, a collagen precursor solution was pipetted around the needle and allowed to gel. Once gelled, the needle was removed, which left a now-hollow channel within the collagen matrix. Using the viscous finger patterning technique^35^, we then lined the lumen of the channel with a basement membrane, finalizing our mammary duct model as seen in Figure 1B.

To mimic the desmoplasia seen in the development of breast cancer, we used different concentrations of collagen to modulate the mechanical and morphological (fiber density and pore size) properties of our mammary duct model. The obtainable shear moduli ranged from 59 ± 1 Pa for the 4 mg/mL to 346 ± 91 Pa for the 8 mg/mL collagen, which encompassed the range that has been measured for healthy and tumor breast tissue ^36^ (Figure 1C). For the remainder of the study, we chose to compare cells grown within models of 4 mg/mL (low) and 8 mg/mL (high) collagen, since they provided the clearest distinction between healthy and diseased tissue properties, respectively.

### Validation of the mammary duct model using phenotypically distinct breast cancer cell lines

We first tested our organotypic mammary duct model by examining its ability to capture the behaviors of two well-established, phenotypically divergent breast cancer cell lines with respect to the two different collagen concentrations. The first, the triple-negative MDA-MB-231 cell line, is known to display an invasive phenotype in 3D and on collagen gels. The second, the luminal-like MCF7 cell line, is known to form clusters in 3D and on collagen gels ^37–39^.

MDA-MB-231 cells within mammary duct models of both low and high collagen concentration displayed little cell-cell interactions (Figure 2A). Instead, they exhibited elongated morphologies with spindle-like shapes that led to an average aspect ratio of 2.91 ± 0.11 and 2.77 ± 0.63 in the low and high collagen concentration models, respectively (Figure 2C, left). The elongation is expected for this highly metastatic cell line, since they are highly motile and minimally express cadherins ^37,40^. This is further seen in the propensity of the MDA-MB-231 cells to invade into the collagen matrix. Counting the total nuclei in cross sections of the mammary duct models (Figure 2B) as well as those that were at least 20 μm (one average cell length) into the matrix from the lumen surface, we calculated the percentage of invasive cells and reported the average invasive distance (Figure 2D). In the lower collagen concentration models, more cells invaded into the surrounding matrix (29.0 ± 9.1% as compared to 2.6 ± 1.8 % in the high collagen concentration), but the distance travelled by the MDA-MB-231 cells was increased in higher collagen concentrations (Figure 2D). Increasing collagen concentration decreased the number of cells that were invasive, but those that did invade traveled a greater distance into the matrix.

**Figure 2.**
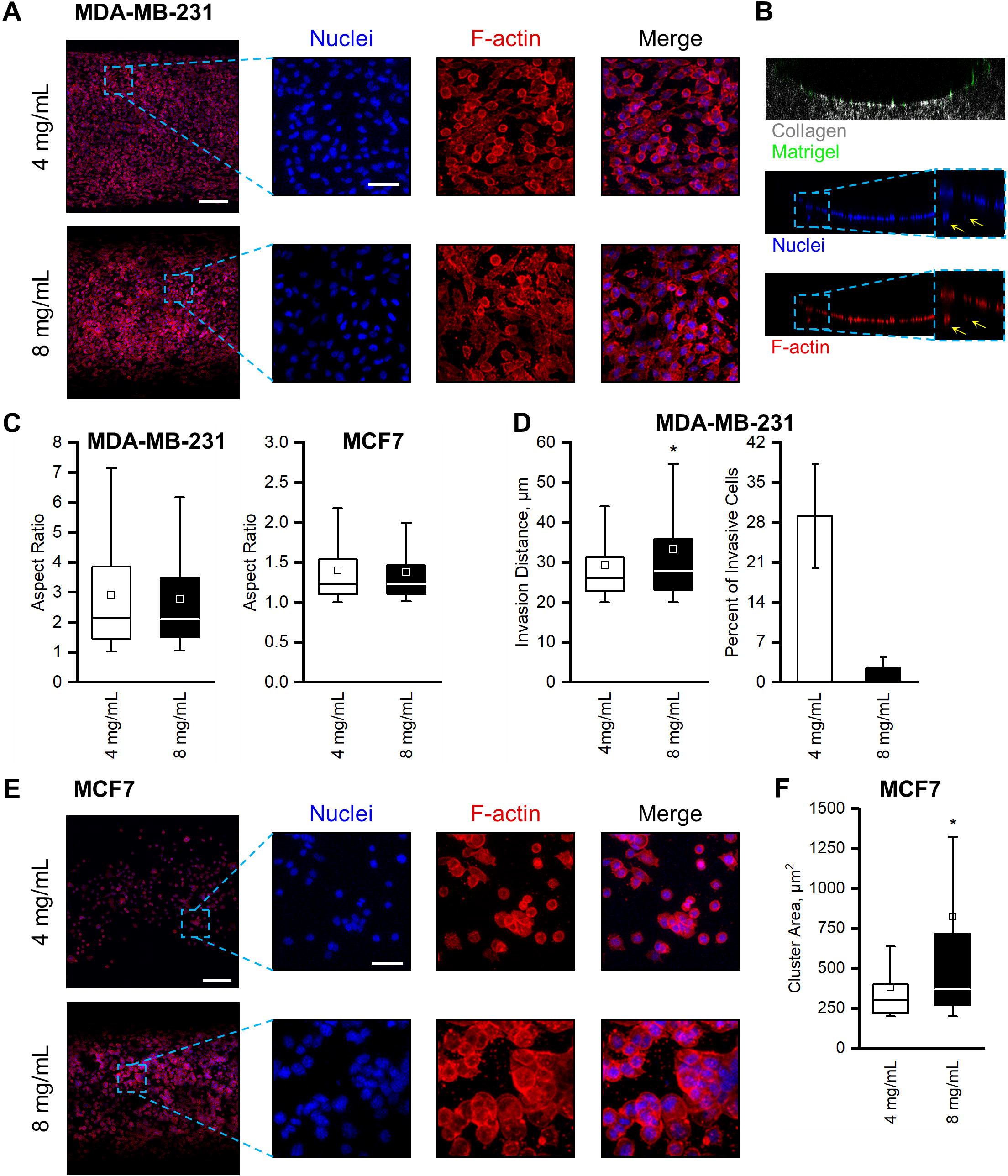
Validation of mammary duct models with MDA-MB-231 and MCF7 cells. A) Representative max projections of MDA-MB-231 cells within mammary duct models of 4 mg/mL (top) and 8 mg/mL (bottom) collagen, cultured for 72 hours. Scale bar is 200 μm. Insets show nuclei (blue), F-actin (red), and a merge of a select region, denoted by the dotted blue lines. Scale bar is 50 μm. B) Representative leftward slice of a cross-section of a mammary duct model at 8 mg/mL showing a basement membrane-lined (green) collagen microchannel (gray) with MDA-MB-231 cells invading into the matrix illustrated by the nuclei (blue) and Factin (red) signals beyond the lumen, denoted by the yellow arrows. C) Measured aspect ratios of MDA-MB-231 (left) and MCF7 (right) cells within mammary duct models of each concentration. D) Percent of invasive MDA-MB-231 cells and their average invasion distance, corresponding to panel B. E) Similar to (A), but with MCF7 cells. F) Characterization of the cluster area of MCF7 cells within the mammary duct models. * denotes p < 0.05.

MCF7 cells, on the other hand, remained rounded and formed clusters in both the low and high collagen concentration mammary duct models (Figure 2E). These morphologies, especially clustering, were expected as MCF7 breast cancer cells highly express cadherins, which drive cohesive interactions and tightly packed aggregates ^37,40^. Few spread or elongated cells were observed; rather, the rounded MCF7 cells had average aspect ratios of 1.40 ± 0.02 and 1.38 ± 0.03 in low and high collagen concentrations, respectively (Figure 2C, right). The clustering behavior, however, was affected by the collagen concentration. MCF7 clusters in the low collagen concentration formed significantly smaller clusters than in the high concentration models (Figure 2F).

By culturing the invasive MDA-MB-231 and cluster-prone MCF7 cells within our organotypic mammary duct model, we validated that our model was able to capture the distinct behaviors associated with the respective breast cancer cell line and that the behaviors were influenced by collagen concentration. These experiments provide evidence that our model can be used to explore breast cancer behaviors within a more physiologically relevant system.

### Morphological characterization of the isogenic MCF10 series and MCF7 breast cell lines within the organotypic mammary duct models

The MCF10A cell line is commonly used as a non-transformed human mammary epithelial cell (MECs) to model normal breast cultures ^11,41,42^. Researchers have modified this cell line through transfections, cloning, and multiple passages to produce the MCF10DCIS.com cell line that can form DCIS comedo-like lesions within mice and the MCF10CA1 cell line that can rapidly and readily form tumors within mice ^41–45^. Together, this MCF10 series encompasses a range of cell lines with a genetically similar background, but with modifications that allow them to model breast cancers at different stages – from healthy to aggressive ^41,42,46^. As we have demonstrated how our organotypic mammary duct model is able to capture distinct behaviors of established breast cancer cell lines and that collagen concentration can impact invasion and clustering, we aimed to study this isogenic series of breast cells within our mammary duct model and compare them to MCF7s, an established DCIS cell line.

Observations on the tissue scale indicated that the organization of cells from the different MCF10 lines on the luminal surface of the mammary duct model were distinct (Figure 3). All MCF10 cell lines adhered to the lumen surface in both collagen concentrations. The healthy MCF10A cells organized themselves, in both collagen concentrations, into a cohesive sheet covering the luminal surface with distinct cell boundaries (Figure 3A). This epithelial-like behavior of a normal MEC line is expected and again affirms the functionality of the model. However, genetic modifications and culture mutations often lead to phenotypes that deviate from epithelial and instead towards more mesenchymal or invasive as evidenced by the MCF10 variants. Thus, our organotypic mammary duct model provides a platform to begin to understand physiological cell-matrix interactions in this breast cancer progression model.

**Figure 3.**
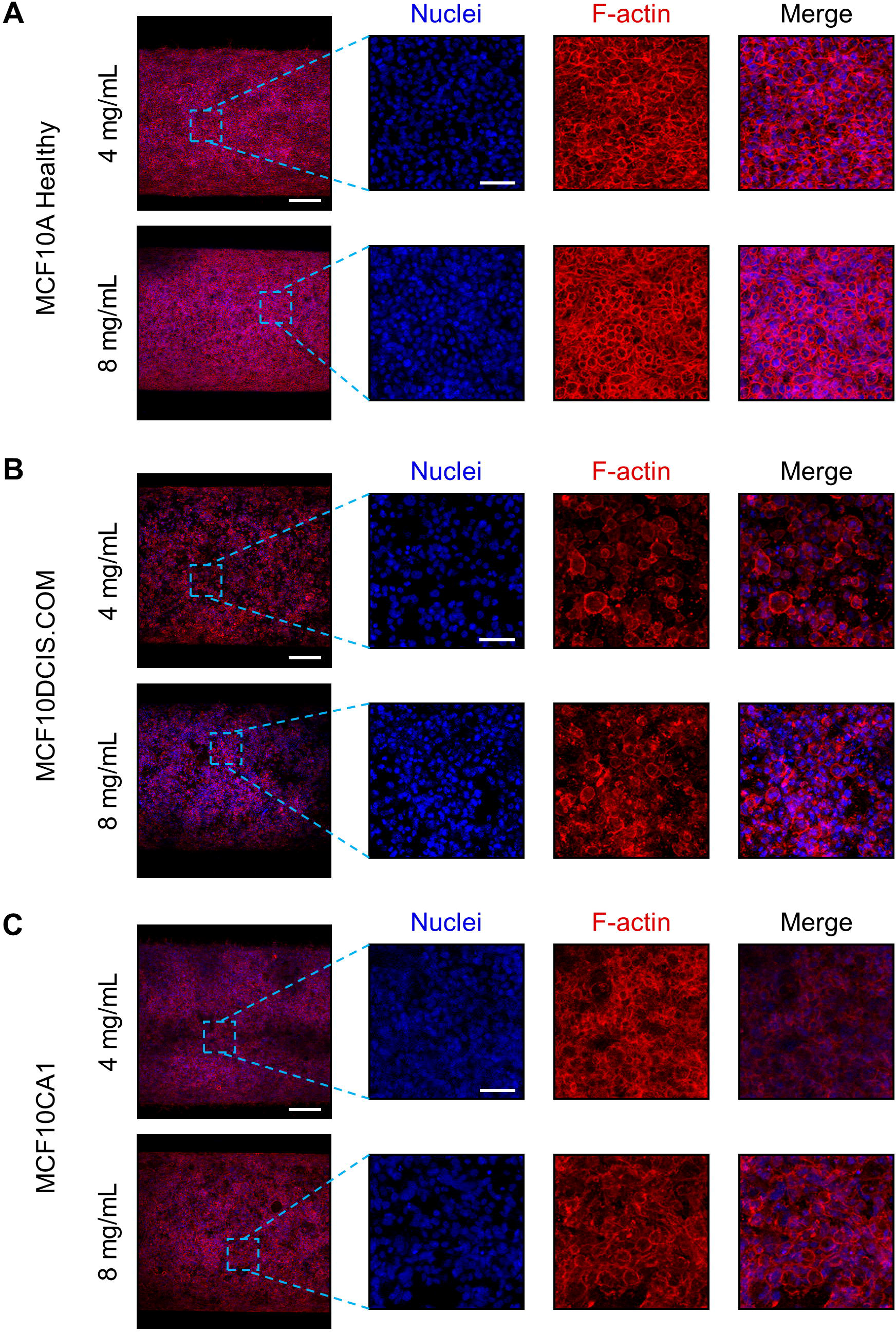
MCF10 series within the mammary duct models. Representative max projections (left, scale bar 200 μm) of cells within mammary duct models of different collagen concentrations grown for one week. Insets (scale bar 50 μm) show nuclei (blue), F-actin (red), and a merge of a select region as denoted by the dotted blue lines. A) MCF10A B) MCF10DCIS.COM C) MCF10CA1.

As expected, the arrangement of the cells of all breast cancer lines on the luminal surface was disorganized and each displayed different degrees and types of aggregation. The MCF10DCIS.COM cells did not evenly cover or form cohesive aggregates to line the luminal surface. Instead, the cells appeared to loosely interact with and form single-cell spots on the luminal surface, which appeared denser at the higher collagen concentration (Figure 3B). The most aggressive MCF10CA1 cells appeared to cover much of the luminal surface, but in a disorganized, web-like network with pockets of space and indistinct cell boundaries (Figure 3C). In contrast, MCF7 cells—now at a higher seeding density and longer culture period—formed tight clusters in both collagen concentrations and significantly increased coverage of the luminal surface at the higher concentration (Figure 4).

**Figure 4.**
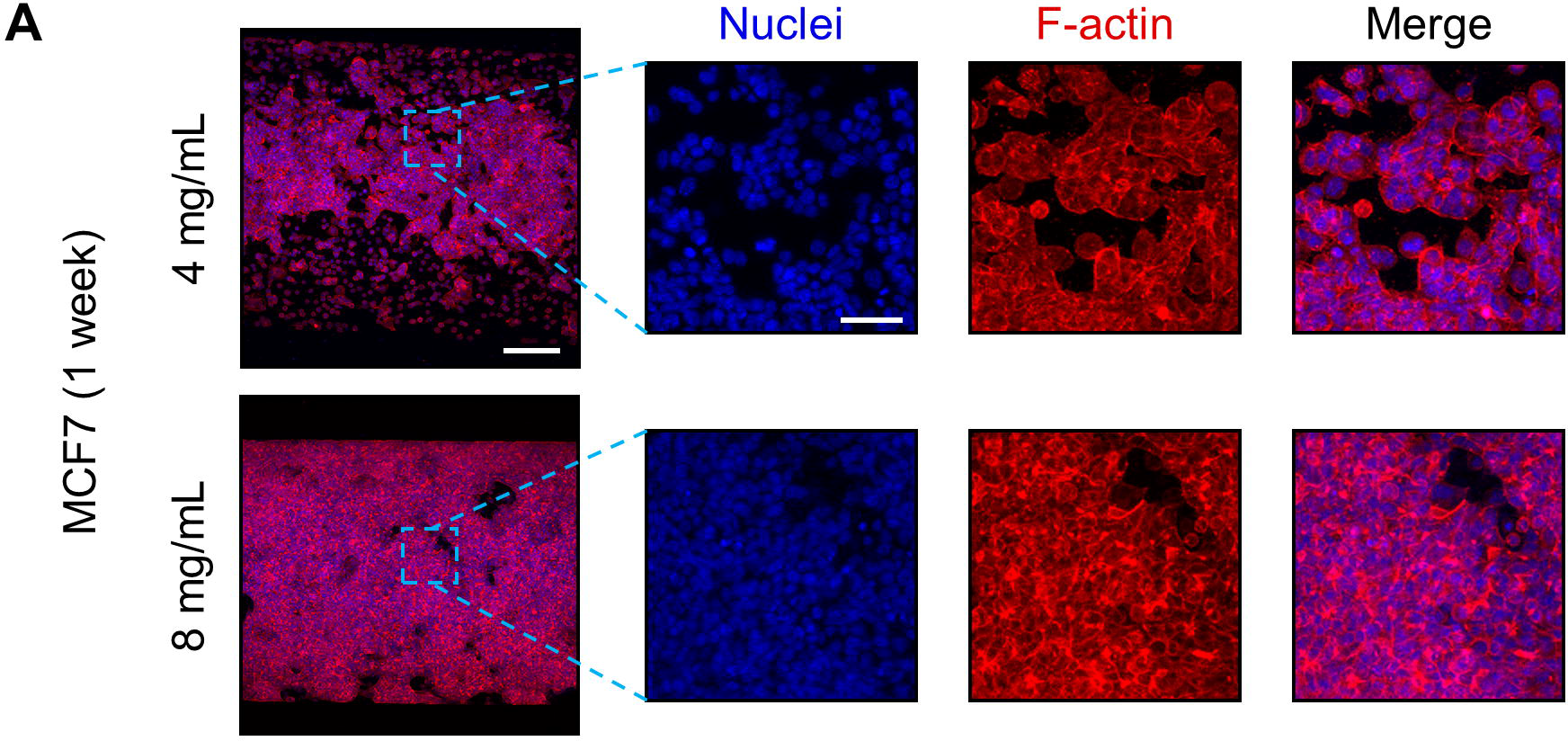
MCF7 cells cultured for 1 week within the mammary duct models. Representative max projections of MCF7 cells within mammary duct models of different collagen concentrations (left) seeded for one week. Scale bar is 200 μm. Insets show nuclei (blue), F-actin (red), and a merge of a select region as denoted by the dotted blue lines. Scale bar is 50 μm.

We began to characterize the organization of the cells on the luminal surface by measuring the distance between neighboring nuclei (Figure 5A) in cross-sectional images of the mammary duct models (Supplementary Figure 2). The MCF10A cells had no difference in nuclei spacing between the two collagen concentrations, indicating that their epithelial-like organization was not disrupted in the range tested. In contrast, the MCF10DCIS and MCF10CA1 cells presented with significantly greater nuclei spacing when cultured within the higher collagen concentration model. The variance of the nuclei spacing was greater, however, in the lower collagen concentration in the case of the MCFDCIS.COM, suggesting more irregularity in the organization of the cells, which is observed in Figure 3B (top row). Our observations of MCF7 cell organization on the luminal surface was also confirmed by this measure. Specifically, the distance between nuclei was larger and had a larger variance at the lower collagen concentration. This correlates with the large spacing seen between aggregates of MCF7 cells on the luminal surface of the lower collagen concentration models, whereas in the higher concentration, the aggregates covered more of the luminal surface (Figure 4).

**Figure 5.**
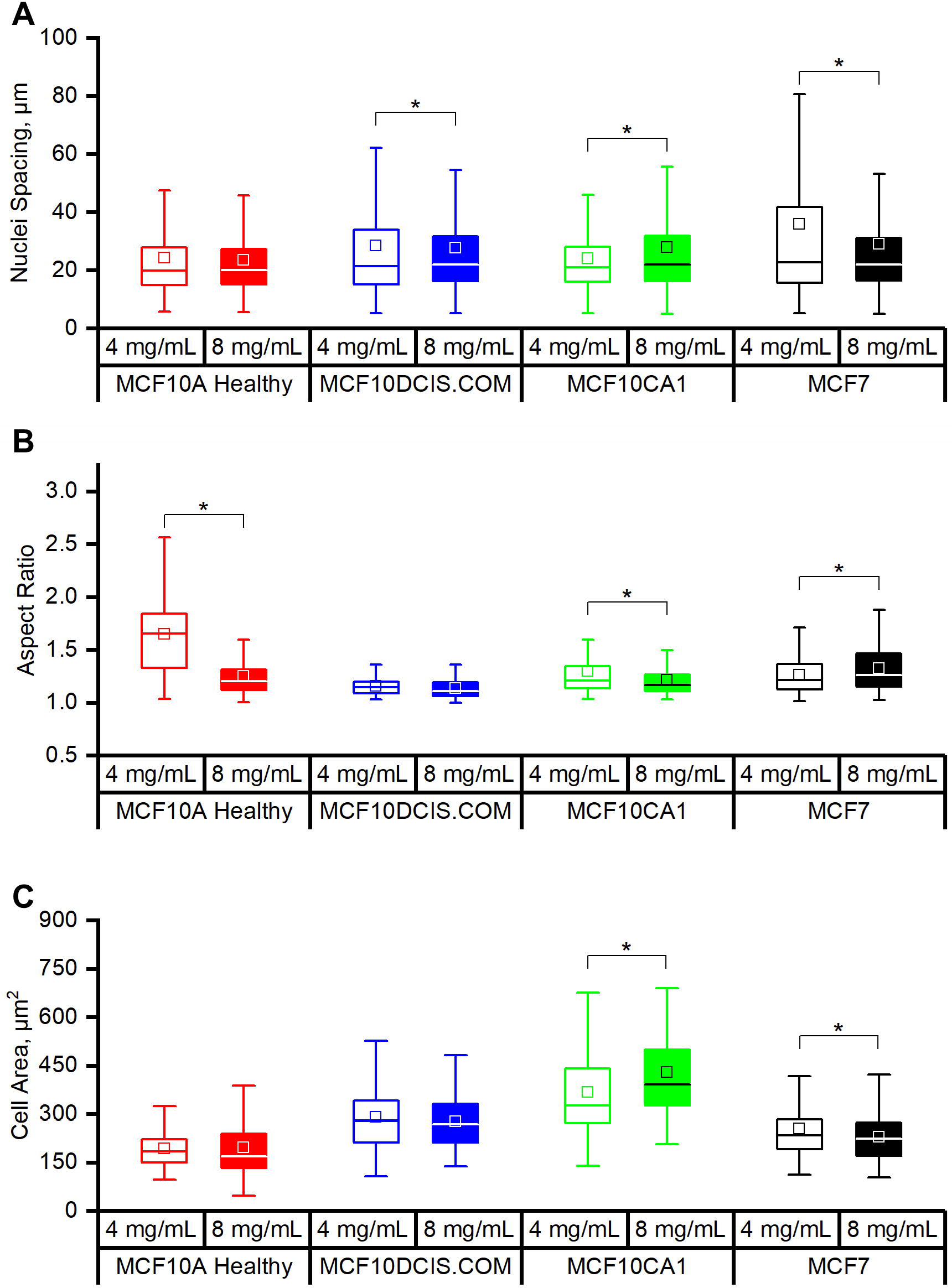
Morphological characterization of cell lines within the mammary duct models. A) Measured distances between neighboring nuclei of breast cells along the lining of mammary duct models. Measured aspect ratios (B) and cell areas (C) of breast cells within mammary duct models of different concentrations. * denotes p < 0.05.

Collagen concentration also influenced the individual cell morphology. Cell spread area and/or aspect ratio was influenced by collagen concentration for all cell lines except the MCF10DCIS.COM (Figures 5B and 5C), which remained rounded with a mid-range spread area. In mammary duct models with the higher collagen concentration, both the MCF10A and MCF10CA1 cells had lower average aspect ratios, but the spread area of the MCF10A cells did not change while the MCF10CA1 did. In contrast, increasing collagen concentration led to an increase in the aspect ratio of the MCF7 cells together with a reduction in their spread area. Combining these measurements with our observations of the tissue scale organization, it appears that the cells that are affected by collagen concentration are those that tend towards some level of organized cohesive interactions between themselves and the luminal surface (i.e., all except MCF10DCIS.COM).

### Breast cell behavior at the mammary duct model luminal surface: hyperplasia and protrusion of breast cells

In analyzing how the different cell lines covered the luminal surface, we recognized that the aggregating behavior and indistinct cell boundaries indicated piling of cells on the lumen. We therefore characterized the thickness of the cell layer for each cell line and collagen concentration. This was done in two ways: i) measuring piling thickness in higher magnification images taken at the mid-height plane of the channel at the luminal surface and ii) by measuring the entire cell-covered area in cross-sections of the bottom half of the lumen (schematics in Supplemental Figure 2).

Representative images of the of the luminal surface at the mid-height plane of the channel are shown in Figure 6A. The healthy MCF10A cells formed a thin, one cell layer along the luminal surface, consistent with their epithelial sheet appearance in Figure 3A. Conversely, all breast cancer cells exhibited piling of the cells that resembled hyperplasia (Figure 6A). The MCF10DCIS and MCF10CA1 each piled on the wall with increasing thickness, correlated with their increasing aggressiveness (Figure 6B). The MCF7 cells produced the thickest cell layers, especially when cultured within the higher collagen concentration model (Figure 6B). This trend towards occlusion of the mammary duct by the cancerous breast cells in part demonstrates their ability to model DCIS.

**Figure 6.**
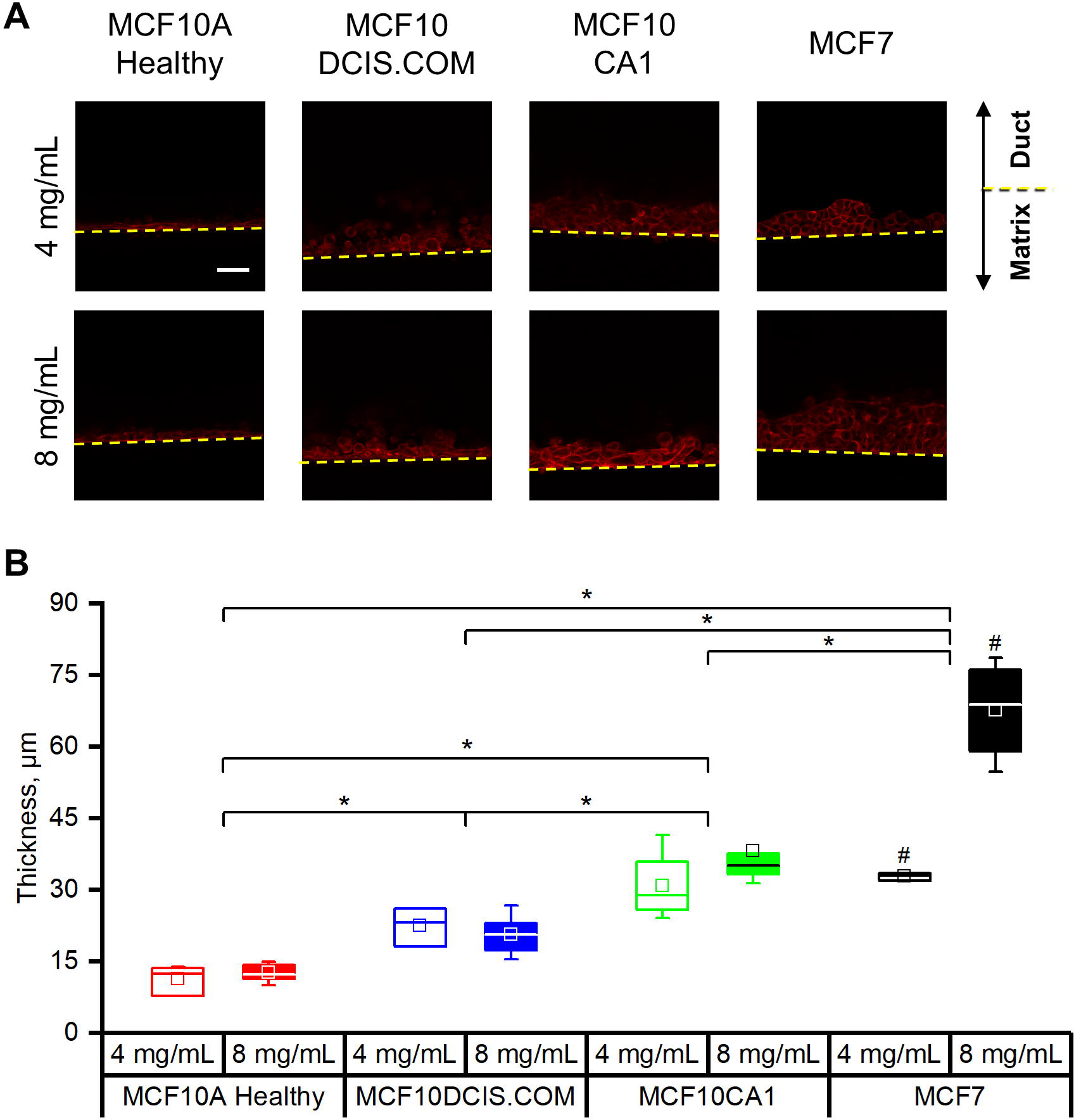
Breast cancer cells can demonstrate hyperplasia within the mammary duct models. A) Representative confocal slices of F-actin from breast cells along the wall of the mammary duct models of 4 mg/mL (top) and 8 mg/mL (bottom) collagen. Dotted yellow lines denote wall of the mammary duct. Upper portions of the images continue into the duct, while the lower portions are the collagen matrix. Scale bar is 50 μm. B) Measured thicknesses of breast cells along the lumen of mammary duct models of different collagen concentrations. * denotes p < 0.05.

The coverage of the luminal surface, according to the projection images through the bottom luminal surface (Figures 4 and 5), was not uniform for most of the cell lines. Therefore, as a complementary measurement to the thickness of the cell piling of Figure 6B, we also measured the percent lumen coverage from cross-sections of the mammary duct models (bottom half of the channel). In each cross-section image, the total area of the F-actin signal was divided by an idealized healthy epithelial sheet layer at the luminal surface: the arc length of the bottom half of the channel multiplied by the average luminal surface-attached cell thickness of the respective cell line. The MCF10DCIS and MCF7 cells exhibited similar trends with an increase in lumen coverage when cultured in the higher collagen concentration model (Figure 7A). This trend is consistent with our previous tissue scale and nuclei spacing observations. The thickness of MCF7 cell piling within the higher collagen concentration model can also be observed with the lumen coverage percentage exceeding 100%. However, observations of the MCF10A and MCF10CA1 cells showed that their lumen coverage similarly exceeded the ideal lumen coverage, though at lower collagen concentrations (Figure 7A). This was striking since the cells were not excessively piling along the lumen (MCF10A, Figures 6A and 6B) nor were the thicknesses at the luminal surface different between the two collagen concentrations (MCF10CA1, Figure 6A and 6B).

**Figure 7.**
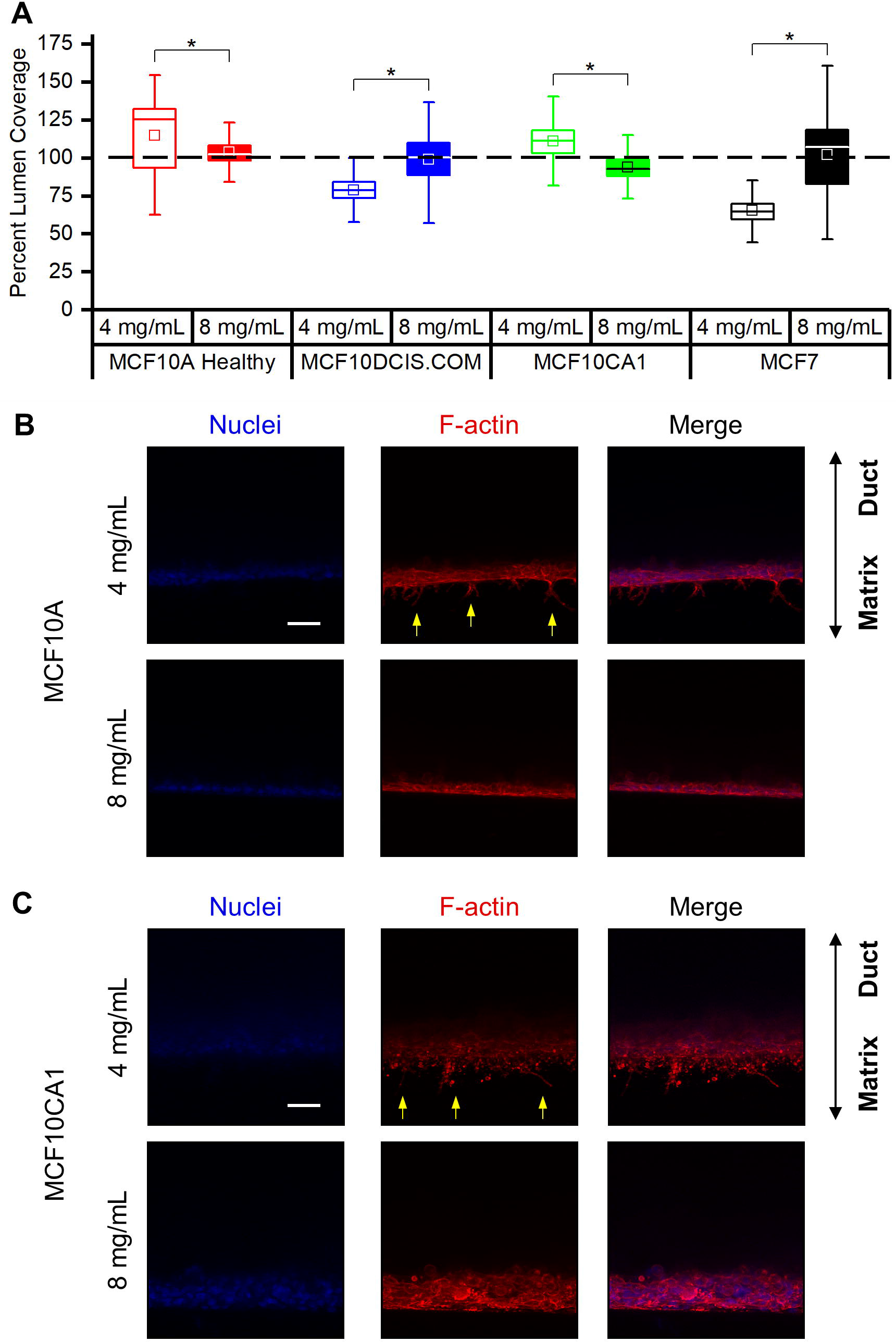
The organotypic mammary duct model can capture the protrusion of breast cells. A) Percent of F-actin coverage of breast cells along the perimeter of mammary duct models. Dotted black line denotes 100% coverage. * denotes p < 0.05. B) Max projections of nuclei (blue, left), F-actin (red, center), and merge (right) of MCF10A cells along the lumen of mammary duct models of different collagen concentrations. Upper portions of the images continue into the duct, while the lower portions are the collagen matrix. Yellow arrows point to F-actin protrusions intro the collagen matrix. C) Same as (B), but with MCF10CA1 cells.

To determine why the MCF10A and MCF10CA1 cell lines had exceeded the ideal lumen coverage percentage, especially at the lower concentration, we returned to the aforementioned mid-height plane images (Figures 7B and 7C), but this time focusing on the matrix side of the lumen-matrix interface. From max projections, we were able to observe protrusions extending from the luminal surface into the matrix by both cell lines at the lower collagen concentration. The fluorescence signal from these protrusions and invasive cells raised the percent lumen coverage measurement for the MCF10A and MCF10CA1 significantly above 100% (Figure 7A).

## DISCUSSION

The need for more physiologically relevant models for breast cancer research remains high as the limited understanding of the transition from DCIS to IDC continues to lead to overdiagnosis and overtreatment. We have developed an organotypic mammary duct model that is mechanically and morphologically tunable to mimic the evolving microenvironment of breast cancers^21–23,47^. Here, we isolate one aspect of the tumor microenvironment—collagen concentration—although the biological and physical complexity of the model could be readily increased (e.g., co-cultures, growth factors, microfluidics, and additional matrix proteins). Furthermore, we recognize that increasing collagen concentration not only increases the mechanical stiffness, but is coupled with increasing ligand density and decreasing pore size ^39,48,49^. Studies expanding on the capabilities of this model should control for such coupled mechanical-morphological effects using methods to alter collagen gelation kinetics and/or cross-linking^22,39,50,51^. Our model also includes a basement membrane layer, which is an important structure as it can act as a barrier to disease progression^52^. We used Matrigel, which, in spite of batch-to-batch variability of components and growth factors, is regularly used as a substrate for studying breast cells in 3D because it is one of the few materials available to mimic a basement membrane ^53^. This model builds upon currently investigated systems—including spheroids in liquid overlays or full embedment into reconstituted basement membrane—that do not consider tissue architecture and other channel models that do not consider changes in matrix physical properties^11,54,55^.

We first validated our model by testing for, and observing, the expected morphologies of two well-established breast cancer cell lines that represent divergent phenotypes: invasiveness (MDA-MB-231) and clustering (MCF7). We also found that varying collagen concentration— our preliminary model of desmoplasia—alters invasive and clustering behaviors. Our findings confirm the dual roles of an activated extracellular matrix as a steric obstacle and a promoter ^56^. Higher collagen concentration is typically associated with denser matrices with smaller pore sizes ^56,57^. The nature of such a matrix to act as an obstacle has been observed in the decreased the number of MDA-MB-231 cells invading into collagen matrices ^39^, which we also observed (Figure 2B). A denser matrix, the result of increasing collagen concentration, can also act as a promoter of invasion distance as has observed for glioma spheroids in collagen I gels of increasing concentration ^58^, similar to our observation of increased invasive distance of MDA-MB-231 cells in the higher collagen concentration. Several factors have been proposed to contribute to promoting cell invasiveness in higher collagen concentrations. These include increased rigidity of the matrix, allowing cells to generate the forces necessary for spreading and migration ^56^ or increase their MMP activity ^59^, and increased collagen fiber alignment that could direct cancer cell protrusion rate, and thereby motility ^57^.

After our validation experiments, we investigated the response of an isogenic breast cancer cell series—spanning the range of healthy to malignant—within the organotypic mammary duct model. To our knowledge, this is the first time the MCF10 series is comparatively cultured within such an organotypic, mammary duct architecture *in vitro* model (i.e., not a liquid overlay, embedment, or xenograft). The combination of our model and the cell line series highlighted the effectiveness of the cell lines to capture phenotypes associated with various stages of breast cancer. All MCF10 cell lines adhered to the lumen surface in both collagen conditions, which was interesting as these cells have also been recently observed to form spherical, acinar structures when seeded in the presence of 2% Matrigel on top of flat, soft (elastic modulus ~ 150 Pa) polyacrylamide gels ^60^. We performed morphological characterizations of the cells, from tissue-scale observations of cell organization to the cell morphology on the luminal surface, and compared them to the well-established MCF7 cell line that is also used for modeling DCIS. The cumulative observations are tabulated in Supplementary Table 1.

At the tissue scale, cell organization was disrupted with disease transformation, as compared to the single-cell thickness, epithelial sheet-like nature of the MCF10A cells. The MCF10DCIS.COM cell line was isolated from lesions that resembled a highly proliferative type of DCIS, characterized in xenograft models by an intact basement membrane and comedo necrosis inside the tumor ^43^. In the mammary duct model, the MCF10DCIS.COM cells were rounded and did not effectively form a layer on the luminal surface, instead appearing spotty at both collagen concentrations (Figure 3B). Their lack of organization or morphological response to the changing collagen concentration may be characteristic of their noninvasive, highly proliferative nature. The MCF10CA1 lines, isolated from xenografts of the HRAS-transformed MCF10AT cell line, are capable of lung metastases in xenograft models ^44^ and are also known to form irregular masses in reconstituted basement membrane, in contrast to the spherical structures of the healthy MCF10A cells ^61^. This malignant cell line covered the luminal surface of the mammary duct models, but in a web-like, less organized manner than MCF10A. Their individual morphology was more elongated and less spread in the lower collagen concentration (Figures 5B and 5C), which is in contrast to the oft-observed increase in spreading and elongation with increasing matrix stiffness. The MCF7 cell line, a luminal-like, cluster-forming model of DCIS ^15,39,62^ expectedly formed clusters in the mammary duct model that covered luminal surface with increasing collage concentration (Figures 4 & 7A). As they expanded onto the luminal surface they also elongated and spread less, which may indicate a competition for space due to aberrant proliferation and organization.

Our observations of hyperplasia and cell invasion both further demonstrate the potential of our model to capture morphological characteristics of breast cancer development and be used as a tool, with tissue architectural context, to study the effects of tumor cell-ECM interactions on the progression. The proclivity of the transformed cells to display signs of hyperplasia by piling into the model lumen increased with their degree of aggressiveness (Figure 6B). The healthy MCF10A cells lined the luminal surface with a one-cell layer. However, the breast cancer cells all formed multicellular thick layers that increased with aggressiveness within the MCF10 series. The MCF7 cell piling was most affected by collagen concentration and produced the thickest layer at the higher collagen concentration. Measurements of luminal surface coverage (Figure 7A) revealed that the MCF10A and MCF10CA1 cells, at lower collagen concentrations, protruded into the collagen matrix. This helped to account for the elongated cell shape of those two lines in the lower collagen concentration models (Figure 5B), which retrospectively can be interpreted as an indicator of invasiveness. Other studies have not reported invasion, or even spreading, of MCF10A cells into collagen-basement membrane gels with this nominal magnitude of elastic modulus (~150 Pa) ^21^. This again highlights the need to control for morphological and mechanical properties of reconstituted ECM hydrogels when studying cell behaviors^39,48,49,57^.

## CONCLUSION

Together, we have shown the capacity of our organotypic mammary duct model to capture distinct aspects of breast cancer progression and how the MCF10 series can be used to model the progression of DCIS *in vitro*. We observed the MCF10DCIS and MCF10CA1 cells to begin occlusion of a mammary duct naturally, without the introduction of added hormones or other cells. Moreover, we demonstrated the ability to capture different protrusive behaviors by the MCF10A, MCF10CA1, and MDA-MB-231 cells in a collagen concentration-dependent fashion. The ability of our physiologically relevant model to capture distinct events of breast cancer progression may support the discovery of novel therapeutic targets that are more translatable into the clinic.

## ACKNOWLEDGEMENTS

We thank the National Science Foundation [NSF MRI 1725984] for the Leica SP8 microscope.

