## Supplementary Figure 1, Supplementary Figure 2, Supplementary Table 1 for "An Organotypic Mammary Duct Model Capturing Distinct Events of DCIS Progression"

**SUPPLEMENTAL**

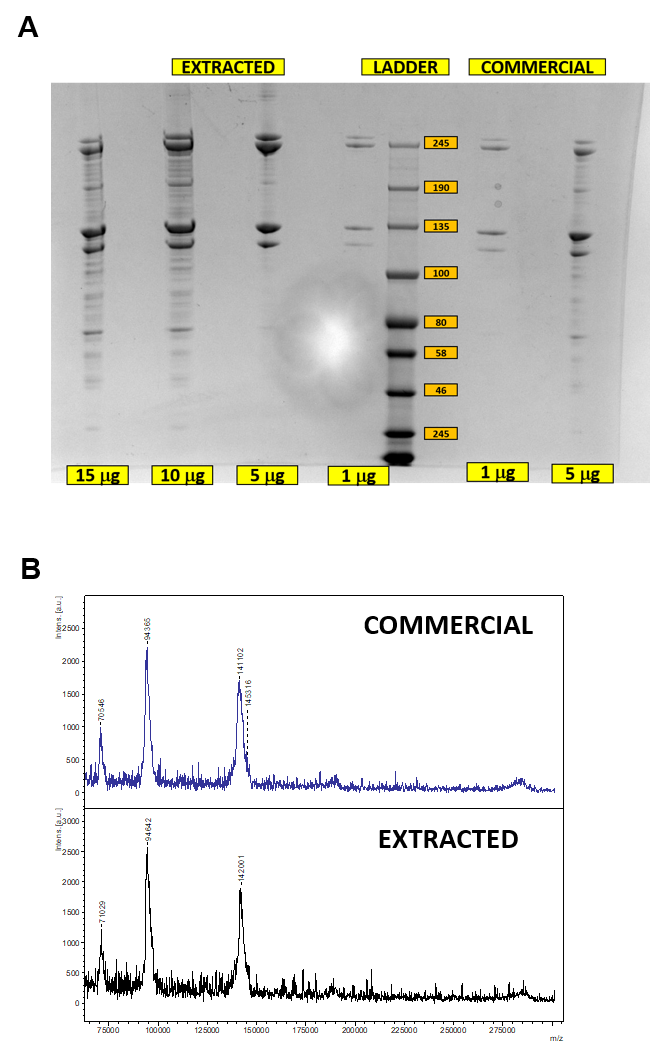

**Supplementary Figure 1. Rat tail collagen purity confirmation.** (A) SDS-PAGE and (B) mass spectroscopy of extracted rat tail collagen type I compared to a commercially purchased item.

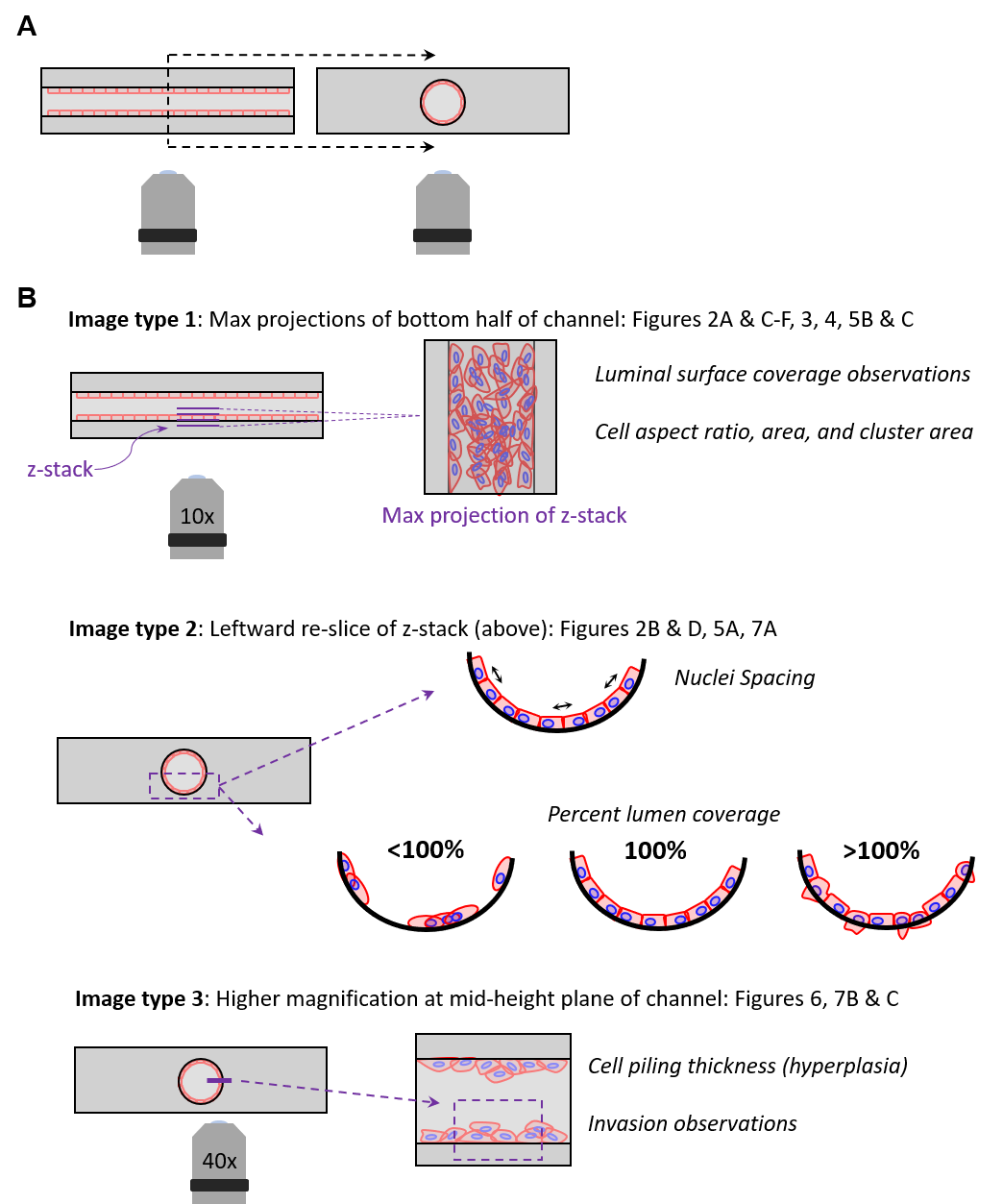
**Supplementary Figure 2: Imaging and image analysis schematics.** A) Positioning of mammary duct sample with respect to objective. B) Diagrams depicting imaging analysis methods for nuclei spacing, actin coverage (percent luminal surface covered), and thickness measurements at the mid-height plane of the channel.

**Supplementary Table 1. Observations of distinct events of breast cancer progression from breast cells within the organotypic mammary duct model.**

| **Cell Line** | **Lumen Organization** | **Hyperplasia** | **Invasiveness** |
| --- | --- | --- | --- |
| MCF10A | Well dispersed and organized in both concentrations. | One cell thick layer in both concentrations. | Protrusions when cultured in lower concentration. |
| MCF10DCIS.COM | Spotty coverage; higher degree of coverage at higher concentration | Noticeable thickness in both concentrations. | None. |
| MCF10CA1 | Web-like spreading in both concentrations. | Noticeable thickness in both concentrations. | Protrusions when cultured in lower concentration. |
| MCF7 | Formation of aggregates. Coverage is highest at higher concentration. | Noticeable thickness; greater at higher concentration. | None. |
